# A reference genome for the long-term kleptoplast-retaining sea slug Elysia crispata morphotype clarki

**DOI:** 10.1101/2023.07.07.548153

**Authors:** Katharine E. Eastman, Amanda L. Pendleton, Mearaj A. Shaikh, Thiti Suttiyut, Raeya Ogas, Paxton Tomko, Gregory Gavelis, Joshua R. Widhalm, Jennifer H. Wisecaver

**Affiliations:** Department of Biochemistry, Purdue University, West Lafayette, Indiana, USA; Purdue Center for Plant Biology, Purdue University, West Lafayette, Indiana, USA; Department of Horticulture and Landscape Architecture, Purdue University, West Lafayette, Indiana, USA

**Keywords:** *Elysia crispata*, Oxford Nanopore Technologies, MinION, long-read assembly, gene expression, comparative genomics, Mollusca, Gastropoda

## Abstract

Several species of sacoglossan sea slugs possess the incredible ability to sequester chloroplasts from the algae they consume. These ‘photosynthetic animals’ incorporate stolen chloroplasts, called kleptoplasts, into the epithelial cells of tubules that extend from their digestive tracts throughout their bodies. The mechanism by which these slugs maintain functioning kleptoplasts in the absence of an algal nuclear genome is unknown. Here, we report a draft genome of the saccoglossan slug *Elysia crispata* morphotype clarki, a morphotype native to the Florida Keys that can retain photosynthetically active kleptoplasts for several months without feeding. We used a combination of Oxford Nanopore Technologies long reads and Illumina short reads to produce a 786 Mbp assembly containing 68,514 predicted protein-coding genes. A phylogenetic analysis found no evidence of horizontal acquisition of genes from algae. We performed gene family and gene expression analyses to identify *E. crispata* genes unique to kleptoplast-containing slugs that were more highly expressed in fed versus unfed developmental life stages. Consistent with analyses in other kleptoplastic slugs, our investigation suggests that genes encoding lectin carbohydrate-binding proteins and those involved in regulation of reactive oxygen species and immunity may play a role in kleptoplast retention. Lastly, we identified four polyketide synthase genes that could potentially encode proteins producing UV- and oxidation-blocking compounds in slug cell membranes. The genome of *E. crispata* is a quality resource that provides potential targets for functional analyses and enables further investigation into the evolution and mechanisms of kleptoplasty in animals.

## INTRODUCTION

The ability to feed on algae and temporarily sequester functional chloroplasts (called kleptoplasts) has convergently evolved multiple times in invertebrates (Van Steenkiste *et al*. 2019). In no lineage is this strategy more common than in the gastropod superorder Sacoglossa, a group of sap-sucking sea slugs (Händeler *et al*. 2009; Maeda *et al*. 2021). In slug digestive cells, phagocytosed algal material is selectively degraded, leaving only kleptoplasts intact, which provide the slug with nitrogen and fixed carbon (Trench *et al*. 1974; Raven *et al*. 2001; Curtis *et al*. 2010; Cruz *et al*. 2020). The length of time kleptoplasts remain photosynthetically active differs between slug species, ranging from a few days in species such as *Elysia cornigera* to over eleven months in *Plakobranchus ocellatus* (Clark *et al*. 1990; Evertsen *et al*. 2007; Händeler *et al*. 2009). Species that can incorporate functional kleptoplasts for up to two weeks of starvation are referred to as short-term-retention forms, and those that retain functional kleptoplasts for over 20 days are considered long-term forms (Christa *et al*. 2014b). Phylogenetic analyses indicate that the ability to retain short-term and long-term kleptoplasts has evolved multiple times in sacoglossan slugs (Christa *et al*. 2015; Maeda *et al*. 2021). Moreover, kleptoplasts acquired from the same algal donor species can have wildly different retention times in different slug species. For example, *Elysia crispata* (synonym *Elysia clarki*) retains long-term kleptoplasts for several months from multiple algal donors, including *Penicillus capitatus*; whereas *Elysia patina* also harbors chloroplasts from *P. capitatus* but the organelles are fully degraded within two weeks (Curtis *et al*. 2010). This suggests that slug-dependent factors contribute to kleptoplast stability and retention. Yet the mechanisms by which saccoglossan sea slugs maintain functional kleptoplasts for extended periods of time remain unclear.

In algae, most chloroplast genes are encoded in the nuclear genome, and key proteins for photosynthesis such as photosystem subunits and light harvesting antennae are synthesized in the cytoplasm and subsequently imported into the chloroplast (Martin and Herrmann 1998; Nowack and Weber 2018). Kleptoplast longevity was, therefore, once hypothesized to stem from horizontal gene transfer (HGT) of algal nuclear-encoded genes to the nuclear genome of kleptoplastic slugs with subsequent targeting of encoded proteins to the kleptoplast (Rumpho *et al*. 2008; Pierce *et al*. 2012). However, transcriptome and genome level analyses of multiple species of kleptoplastic sacoglossans have found no evidence of horizontal gene transfer from algae (Wägele *et al*. 2011; Bhattacharya *et al*. 2013; Maeda *et al*. 2021). Similarly, it was hypothesized that kleptoplastic slugs may translate proteins from retained algal mRNAs to control turnover, stability, and/or repair damaged kleptoplasts (Pierce and Curtis 2012), but a recent transcriptomic analysis of *P. ocellatus* found no evidence to support this “transferome” hypothesis (Maeda *et al*. 2021).

Combined, the lack of evidence for horizontally acquired algal genes or transcripts suggests that evolved nuclear-encoded slug proteins play a role in prolonged kleptoplast retention. For example, kleptoplastic slugs produce propionate pyrones that may function as “sunscreens” in cell membranes (Powell *et al*. 2018). These compounds are *de novo* synthesized from carbon dioxide by slugs in light (Ireland and Scheuer 1979) and are predicted to help limit the formation of damaging reactive oxygen species (ROS) produced by excessive light absorption. Recently, Torres *et al*. (2020) discovered an *Elysia chlorotica* polyketide synthase (EcPKS1) and showed *in vitro* that it can produce propionate pyrone precursors from methylmalonyl-CoA. The authors also found that less UV-reactive 5-propionate products occur in non-photosynthetic and short-term kleptoplastic slugs while longer 7- and 8-propionate pyrones are restricted to slugs performing long-term kleptoplasty, which suggests that novel PKSs have evolved in slugs capable of long-term kleptoplasty.

Genome assemblies are available for two long-term-retention species, *E. chlorotica* and *P. ocellatus*, as well as one short-term species, *Elysia marginata* (Cai *et al*. 2019; Maeda *et al*. 2021). To gain a better understanding of the evolution and underlying mechanisms of kleptoplasty in saccoglossan sea slugs, genomic data across additional species are necessary. Here, we report a high-quality draft genome assembly of *E. crispata* morphotype clarki (Figure 1A,B), a kleptoplastic sea slug native to mangrove swamps and waterways of the Florida Keys (Krug *et al*. 2016). *E. crispata* can retain long-term functional kleptoplasts for three months or more (Middlebrooks *et al*. 2012). Like prior assessments of HGT in kleptoplastic slugs (Wägele *et al*. 2011; Bhattacharya *et al*. 2013; Maeda *et al*. 2021), we found no evidence of algal-derived genes in the *E. crispata* nuclear genome. Rather, our analysis suggests native slug genes involved in regulation of ROS and adaptive immunity may be involved in prolonging the life of kleptoplasts. We performed a phylogenetic analysis of the recently discovered PKS genes in kleptoplastic slugs (Torres *et al*. 2020), identifying four PKS genes in *E. crispata* that may play a role in protecting kleptoplasts from UV and oxidative damage. Taken together, the *E. crispata* genome offers a quality resource for exploring the evolution and mechanisms of long-term kleptoplast retention in Sacoglossa.

**Figure 1.**
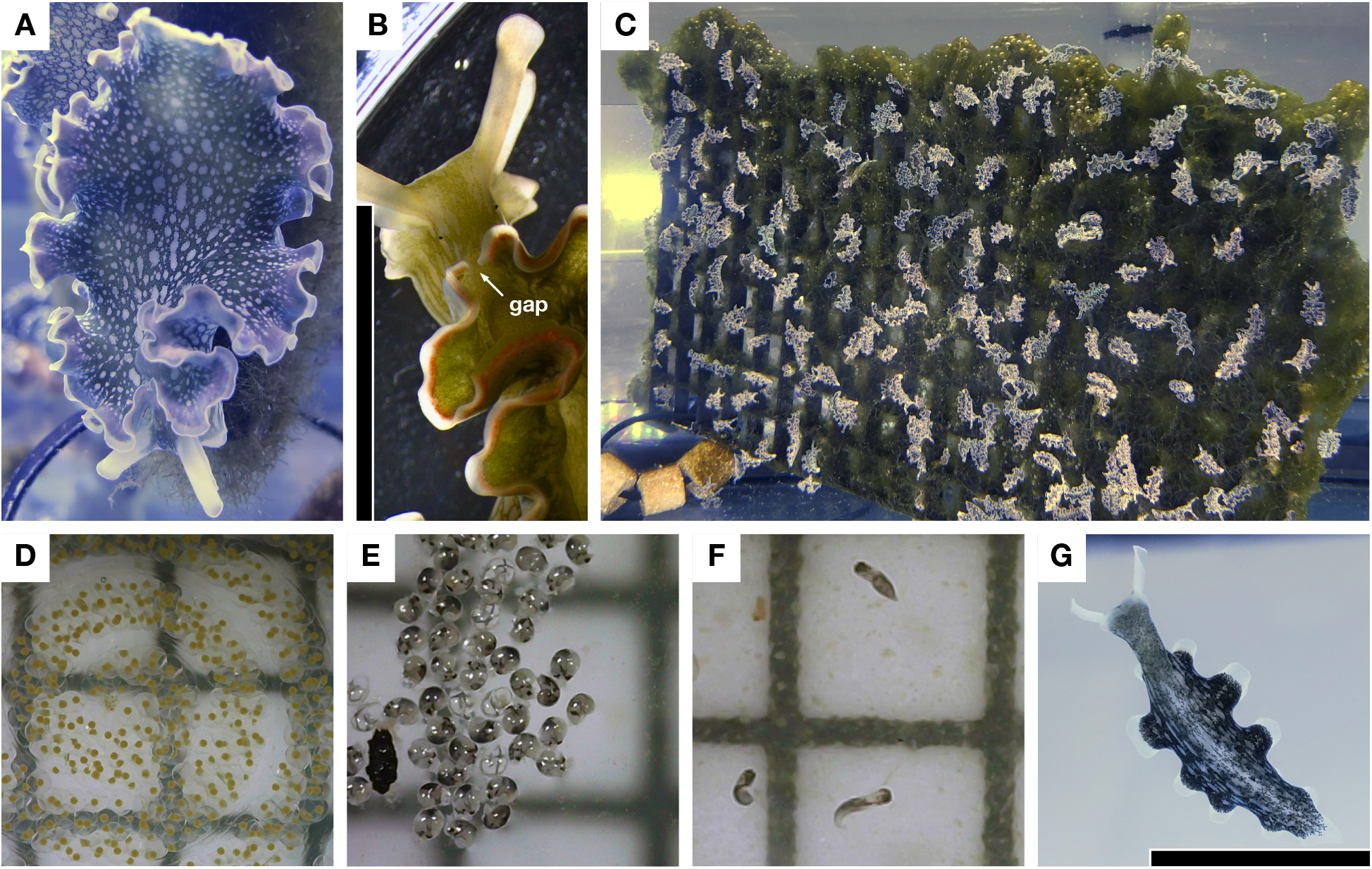
*Elysia crispata* morphotype clarki. A) Wild-caught adult slug, dorsal view, showing near-uniform green coloration with small round white spots indicative of clarki morphotype. B) Close up of wild-caught adult slug showing parapodial gap indicative of clarki morphotype. C) Lab-reared cornucopia of slugs. D) Egg clutch 24 h post deposition. E) Free-swimming veliger. larvae. F) Crawling larvae. G) Lab-reared juvenile slug, ventral view, showing transparent and tapered foot indicative of clarki morphotype; green pigmentation indicates presence of kleptoplasts. Scale bar in B,G = 10 mm. Scale grid in D,E,F = 2 x 2 mm.

## MATERIALS AND METHODS

### Slug and algal material and laboratory culturing in aquaria

Wild-caught *E. crispata* were purchased from a local aquarium shop in Lafayette, Indiana (United States). The slugs were shipped overnight to the shop after being collected from the vertical walls of an undisclosed canal in the Florida Keys (United States).

Slugs were acclimated in 20 L aquaria containing artificial seawater (ASW) prepared using distilled water and Instant Ocean® Reef Crystals at a specific gravity of 1.023 at 25℃ (77℉). Slugs were then transferred to connected 180-200 L aquaria containing ASW recirculated through a 280 L sump tank and a Pentair SMART UV sterilizer to reduce microorganismal growth. The water temperature was maintained at 25℃ using a standard aquarium heater. Aquaria were illuminated with white LED lights (Fluval Sea Marine 3.0 LED Aquarium Lights set at 75% of maximum intensity) on a 12:12 light:dark cycle. Every week, approximately 50% of the system volume was replaced with fresh ASW, and freshly cultured algae was introduced. Aquaria were monitored daily for the deposition of egg masses. Egg masses were collected within one day of deposition.

Macroalgae were cultured on 15 x 30 x 1 cm gridded plastic racks in a separate 220 L aquarium containing recirculated ASW filtered through a 20 L sump tank. To help promote algal growth, a constant turbulent water flow was maintained using three circulation pumps positioned across the top, middle, and bottom of the aquarium in alternating directions. Weekly maintenance consisted of treating the water with 50 µg mL^-1^ rifampicin and changing out 100% of the ASW 24 hours later. Aquarium ASW was supplemented three times per week with one hour of gentle bubbling of compressed carbon dioxide and dosing with 100 mL each of 20 g L^-1^ potassium nitrate (ThermoFisher, Waltham, MA, USA), 1.5 g L^-1^ potassium dihydrogen phosphate (ThermoFisher, Waltham, MA, USA), and 1:20 diluted Guillard’s F/2 trace elements (Algae Research Supply, San Diego, CA, USA).

### Sampling, RNA extraction, and RNA sequencing and processing

Wild-caught adult slugs deposited egg masses approximately 2-3 weeks after being introduced in the laboratory aquaria. Egg samples (Figure 1D) for RNA extraction were rinsed in fresh ASW, flash-frozen in liquid nitrogen, and stored at -80℃ within 24 hours of deposition. Each biological replicate was derived from using the entire egg mass deposited by an individual slug.

To obtain free-swimming veligers, crawling larvae, and juvenile slugs, each remaining collected and rinsed egg mass was transferred to an individual petri dish containing 25 mL of freshly prepared ASW and supplemented with 5 µg mL^-1^ rifampicin and 500 ng mL^-1^ ivermectin to eliminate the potential growth of cyanobacteria, bacterial pathogens, and predators. Individual egg masses were maintained at room temperature under white LED light (Fluval Sea Marine 3.0 LED Aquarium Lights set at 75% of maximum intensity) on a 12:12 light:dark cycle. Egg masses were carefully rinsed and filtered through 52-micron nylon mesh and transferred to new ASW with rifampicin and ivermectin every two days. Free-swimming veliger larvae (Figure 1E) hatched approximately 21-28 d after egg deposition and were collected by centrifuging the ASW media at 1000 x *g* for 1 min. Pellets of free-swimming veliger samples were rinsed in ASW, flash-frozen in liquid nitrogen, and stored at -80℃ until RNA extraction. Each biological replicate was derived from using all the free-swimming veligers that hatched from a single egg mass deposited by an individual slug. The hatchlings from the remaining unused egg masses were allowed to metamorphose into larvae. Crawling larvae (Figure 1F) were maintained as pools derived from individual egg masses and kept in petri dishes as described for egg masses. Crawling larvae samples were prepared as described for veliger samples approximately 3-5 d after metamorphosis. Each biological replicate was derived from using all the larvae derived from a single egg mass. The larvae derived from the remaining unused egg masses were introduced into 16 x 20 cm Pyrex dishes containing freshly prepared ASW, 5 µg mL^-1^ rifampicin, 500 ng mL^-1^ ivermectin, and approximately 1 g of macroalgae. After reaching a length of approximately 1-2 cm, juvenile slugs (Figure 1C,G) were collected, flash-frozen in liquid nitrogen, and stored at -80℃ until RNA extraction. Each biological replicate consisted of three individual slugs.

Total RNA was isolated from 100 mg of pulverized flash-frozen *E. crispata* eggs, free-swimming veligers, crawling larvae, or juvenile slugs using Trizol reagent (ThermoFisher Scientific, Waltham, MA, USA) according to the manufacturer’s protocol. Extracted RNA samples were digested with DNaseI (New England Biolabs, Ipswich, MA, USA) and column purified using an RNA concentrator and cleanup system (Zymo Research, Irvine, CA, USA). The eluted RNA samples were quantified using a NanoDrop spectrophotometer. Purified samples were measured for RNA integrity (RIN) scores at the Purdue Genomics Core facility. Samples with RNA integrity scores ≥ 7 were selected for library prep. Three biological replicates of each *E. crispata* developmental stage of the slug were sent to Novogene Corporation Inc. (Sacramento, CA, USA) for library construction (NEBNext Ultra RNA Library Prep Kit, New England Biolabs) from 1 µg total RNA and Illumina sequencing. Paired-end (2×150 bp) reads were quality filtered and adapter trimmed using fastp v0.20.1 (Chen *et al*. 2018).

### Sampling, DNA extraction, and genome sequencing and assembly

Genomic DNA (gDNA) for Illumina sequencing was extracted from *E. crispata* eggs reared in petri dishes (without algae). Samples were pulverized in liquid nitrogen using a Tissue Lyser II (Qiagen, Hilden, Germany). DNA was extracted using one of two methods, a CTAB-based protocol for samples with high mucopolysaccharide content (dx.doi.org/10.17504/protocols.io.kxygxp81zl8j/v1) modified from (Winnepenninckx *et al*. 1993) or a DNAzol Reagent (Invitrogen, Waltham, MA, USA) protocol (dx.doi.org/10.17504/protocols.io.n92ldprrnl5b/v1). Quantity and quality of DNA for each sample was assessed using a Qubit 4 fluorometer (Invitrogen, Waltham, MA, USA) and a TapeStation 4150 (Agilent, Santa Clara, CA, USA). No difference in quality between the CTAB and DNAzol-based protocols were observed, and the samples were pooled prior to Illumina sequencing. The sequencing library was constructed using an NEBNext DNA library prep kit (New England Biolabs), and 2×150 bp paired-end reads were sequenced using an Illumina NovaSeq 6000 by Novogene Corporation Inc. (Sacramento, CA, USA). Illumina gDNA read quality was assessed by FastQC v0.10.0 (Babraham Bioinformatics 2011). Illumina TruSeq adapters and low quality reads were removed with fastp v0.20.1 (Chen *et al*. 2018) using default parameters.

Illumina reads derived from suspected bacterial contamination were identified using Blobtools v1.1.1 (Laetsch and Blaxter 2017). A first-pass Illumina-only genome assembly was performed by Abyss v2.2.4 (Simpson *et al*. 2009) using a k-mer size of 96. Illumina gDNA reads were aligned to the Abyss assembly using BWA-MEM v0.7.15 (Li and Durbin 2009) to generate a coverage BAM file. Abyss contigs were queried against the NCBI nucleotide (nt) database (accessed September 11, 2021) using blastn v2.11.0 (Camacho *et al*. 2009). DIAMOND v2.0.8.146 (Buchfink *et al*. 2015) was used to query Abyss contigs against a custom protein databases that consisted of NCBI RefSeq release 207 (O’Leary *et al*. 2016) sequences supplemented with additional predicted protein sequences from MMETSP (Keeling *et al*. 2014) and the 1000 Plants transcriptome sequencing project (1KP) (Matasci *et al*. 2014). The custom protein database used in the BlobTools analysis is available from the authors as well as through the following link: https://www.datadepot.rcac.purdue.edu/jwisecav/custom-refseq/2021-08-02/. The BlobTools taxrule ‘bestsumorder’ determined the taxonomic assignment of each contig, prioritizing information from protein hits first. Contigs denoted as non-eukaryotic in origin were flagged, and BBSplit v38.87 (Bushnell 2017) was used to exclude Illumina reads that mapped to these contigs. This step removed 3.4% of reads, as most contigs were either taxonomically unassigned or determined to be of eukaryotic origin (Figure S1). Estimates of genome size and heterogeneity were calculated using GenomeScope2.0 (Ranallo-Benavidez *et al*. 2020).

Genomic DNA for Oxford Nanopore Technologies (ONT) sequencing was extracted from two *E. crispata* sample types: brains dissected from wild adults and whole bodies of wild adults starved for a minimum of 2 weeks (Table S1). Both the CTAB and DNAzol protocols referenced above were used to generate high molecular weight DNA for ONT long-read sequencing (Table S1). Short DNA fragments less than 10 kbp were depleted using an SRE XS kit (Circulomics, Baltimore, MD, USA) according to the manufacturer’s protocol. At least 1.9 μg of high molecular weight genomic DNA was used as input for ONT LSK-109 library ligation kits and sequenced on R9 MinION flow cells. Six flow cells were necessary to generate sufficient read depth for whole genome assembly (Table S1). Base calling of ONT reads was performed with Guppy v6.1.2 (Oxford Nanopore Technologies, Oxford, UK). ONT reads were quality filtered by Filtlong (Wick 2022) using the Illumina reads as an external reference and with the following parameter settings: --trim --split 500 --keep_percent 90 --min_length 3000.

Two programs for genome assembly were evaluated: CANU v2.2 (Koren *et al*. 2017) using an estimated genome size of 711 Mbp and Flye v2.9 (Kolmogorov *et al*. 2019) using the -- keep-haplotypes and --nano-raw parameter settings. Three programs for error correction and genome polishing were evaluated: Racon v1.5.0 (Vaser *et al*. 2017), Medaka v1.6.0 (nanoporetech 2022), and NextPolish v1.4.0 (Hu *et al*. 2020). Racon and Medaka polishing runs were performed using ONT reads only. NextPolish polishing runs were performed using ONT and Illumina reads as references. Racon polishing was iteratively performed up to four times. All polishing programs were run using default parameters. Polished assemblies were compared to each other and to the unpolished starting assemblies using QUAST-LG v5.2.0 (Mikheenko *et al*. 2018) and BUSCO v4.0.6 (Manni *et al*. 2021) run in genome mode with the eukaryote_odb10, metazoan_odb10, and mollusca_odb10 lineage datasets. The Flye assembly was selected as the final published assembly as it resulted in longer contigs (Table S2) and better BUSCO recovery compared to CANU (Table S3). Polishing results were comparable across the different iterations. No single iteration performed best for all BUSCO odb10 datasets: three rounds of Racon polishing yielded the best BUSCO recovery of the eukaryote_odb10 dataset (96.08% complete and single copy); one round of Racon polishing followed by Medaka and NextPolish yielded the best BUSCO recovery of the metazoan_odb10 dataset (93.82% complete and single copy); and a single round of Medaka polishing yielded the best BUSCO recovery of the mollusca_odb10 dataset (87.76% complete and single copy). Therefore, the three best iterations were all carried forward to gene annotation and protein prediction (see Genome and gene annotation methods section below). Ultimately, one round of Racon polishing followed by Medaka and NextPolish was selected as the final polishing strategy based on its superior BUSCO recovery of all three odb10 datasets when run in protein mode (Table S3).

### Genome and gene annotation

*De novo* repeat identification was performed using RepeatModeler v2.0.1 (Flynn *et al*. 2020), and the resulting repeat library was used to inform repeat masking with RepeatMasker v4.0.7 (Smit *et al*. 2017). Gene model and protein prediction was conducted using BRAKER2 v2.1.5 (Hoff *et al*. 2019; Brůna *et al*. 2021). For the initial run, BRAKER2 was supplied the *E. crispata* assembly with 1) repeats soft-masked, 2) a custom protein database comprised of metazoa_odb10 and the predicted proteomes of 23 mollusk genome assemblies on NCBI (Table S4), and 3) the *E. crispata* Illumina RNA-Seq data aligned to the genome using STAR v2.7.10 (Dobin *et al*. 2013). We noticed that highly conserved, multi-copy genes (*e.g.*, tandem duplicates of histone protein H3) were missing from this initial run, and manual investigation revealed that these types of gene families had been incorrectly modeled as repetitive elements by RepeatModeler. Therefore, we performed a second BRAKER2 run using the unmasked assembly. Gene models resulting from BRAKER2 annotation with and without the use of the masked reference assembly were independently filtered to retain the highest quality gene models. Unique gene models resulting from the unmasked run were identified through coordinate intersection with the masked gene set using BEDTools (Quinlan and Hall 2010) and were added to the final gene set if they satisfied the following criteria: 1) had a significant BLASTp hit (e-value < 1e^-3^) to a database of model animal proteomes that included human (GRCh38), zebrafish (GRCz11), mouse (GRCm39), and two mollusks *Crassostrea virginica* (C_virginica-3.0) and *Pomacea canaliculata* (v1.0); 2) did not intersect with a classified repeat element (LINE, SINE, LTR, etc.); and 3) did not have exons that were > 50% simple sequence repeats. A flowchart of these curation steps is provided in Figure S2.

To identify mitochondria- and kleptoplast-derived contigs, predicted proteins from the *P. ocellatus* mitochondrial genome (NCBI accn: AP014544.1) and *R. lewmanomontiae* (order: Bryopsidales) chloroplast genome (NCBI accn: AP014542.1) were queried using tblastn v2.11.0 (Camacho *et al*. 2009) against the *E. crispata* genome assembly. Contigs with two or more significant hits (e-value ≤ 1e^-5^) to these organellar proteins were annotated using GeSeq (Tillich *et al*. 2017), and OGDRAW (Greiner *et al*. 2019) was used to generate a physical map of each contig. The physical maps were manually inspected, and one contig was flagged as mitochondrial in origin (Figure S3). An additional nine contigs were flagged as kleptoplast or kleptoplast fragments (Figure S4). All organelle-derived contigs were excluded from the nuclear genome assembly and can be downloaded from FigShare (see Data Availability).

During the Alien Index analysis (see Methods section on HGT below), we flagged 194 contigs as bacterial contamination, which were excluded from the nuclear genome assembly. Gene models were removed if they were located on any contig that was flagged from the Alien Index pipeline. Additionally, we flagged the proximal end of contig_7836 as misassembly likely resulting from kleptoplast contamination. The region consisted of 16,750 kbp and contained four genes, including two genes annotated as Photosystem I P700 chlorophyll a apoprotein (PsaA/PsaB)-like. Typically, *PsaA* and *PsaB* are both encoded on the chloroplast genome of chlorophyte algae. To investigate the accuracy of the assembly at this locus, Illumina gDNA and ONT reads were aligned to the genome using BWA MEM (Li and Durbin 2009) and minimap2 (Li 2016) using default parameters, respectively. Alignments relative to genes and repeat elements were visualized using Integrated Genomics Viewer (Robinson *et al*. 2011). No Illumina reads joined the locus in question to the rest of contig_7836. Only a single ONT read mapped to the “gap” between the contig ends, but it mapped poorly (Figure S5). Altogether, this indicated that this region was likely incorporated into the nuclear genome assembly in error rather than a true HGT event. Therefore, this region of contig_7836 was manually removed from the final genome assembly and the four gene models removed from the final gene set. The resulting *E. crispata* assembly and gene annotations following these steps were designated as final (v1).

### Search for HGT

We searched the *E. crispata* genome for possible HGT using the Alien Index score as previously described (Wisecaver *et al*. 2016). Briefly, each predicted protein sequence was queried against the same custom protein database used for BlobTools (see above) with DIAMOND v2.0.8.146 (Buchfink *et al*. 2015). A custom python script sorted the DIAMOND results based on the normalized bitscore (*nbs*), where *nbs* was calculated as the bitscore of the single best-scoring HSP to the subject sequence divided by the best bitscore possible for the query sequence (i.e., the bitscore of the query aligned to itself). The Alien Index (AI) score is given by the formula: *AI* = *nbsO* − *nbsM*, where *nbsO* is the normalized bit score of the best hit to a species outside of the Metazoa lineage (NCBI:txid33208), *nbsM* is the normalized bit score of the best hit to a species within Metazoa skipping all hits to Placobranchoidea (NCBI:txid71491), a sublineage within Sacoglossa that contains all currently available genomes from kleptoplastic slugs including *E. crispata*. Alien Index scores range from -1 to 1, being greater than 0 if the predicted protein sequence had a better hit to a non-metazoan sequence, suggestive of either HGT or contamination (Wisecaver *et al*. 2016). Contigs were flagged as contamination if the minimum Alien Index score for all genes on a contig was > 0 or if at least half of genes on a contig had an Alien Index score > 0.1. All contamination-flagged contigs were bacterial in origin and were excluded from the nuclear genome assembly (see Methods section on genome and gene annotation above).

We filtered HGT candidates (Alien Index > 0) to those that were most likely to be phylogenetically informative by requiring Alien Index > 0.1 and total database hits ≥ 50. Phylogenetic trees of protein sequences were constructed for all filtered Alien Index-flagged HGT candidates. Full-length proteins corresponding to the top 200 hits (e-value < 1e^-10^) to each HGT candidate were extracted from the local database using esl-sfetch (Eddy 2009). Protein sequences were aligned with MAFFT v7.471 using the E-INS-i strategy and the BLOSUM30 amino acid scoring matrix (Katoh and Standley 2013) and trimmed with trimAL v1.4.rev15 using its gappyout strategy (Capella-Gutierrez *et al*. 2009). Phylogenies were inferred using maximum likelihood as implemented in IQ-TREE v1.6.12 (Nguyen *et al*. 2015) using an empirically determined substitution model and 1000 rapid bootstrap replications. The phylogenies were midpoint rooted and branches with local support < 95 were collapsed using the ape and phangorn R packages (Paradis *et al*. 2004; Schliep 2011). Phylogenies were visualized using ETE v3 (Huerta-Cepas *et al*. 2016) and inspected manually to identify phylogenetically supported HGT candidate proteins.

### Species phylogenies

Sacoglossan 28S ribosomal DNA and histone H3 sequences were downloaded from the NCBI nucleotide database (Figure S6), and corresponding 28S and H3 sequences in *E. crispata* were identified using BLAST. For the 28S phylogeny, sequences were aligned with MAFFT v7.471 using the G-INSI-i iterative refinement method (Katoh and Standley 2013). H3 sequences were aligned with MAFFT using the RevTrans online web portal (Wernersson and Pedersen 2003). Maximum-likelihood phylogenies were constructed using IQ-TREE v2.2.0 (Minh *et al*. 2020) using the built-in ModelFinder to determine the best-fit nucleic acid substitution model (Kalyaanamoorthy *et al*. 2017) and 1000 ultrafast bootstrap replicates. The concatenated ML tree was constructed based on the combined 28S and H3 data matrix (Chernomor *et al*. 2016) consisting of two partitions and 2006 sites. Phylogenies were visualized using ETE v3 (Huerta-Cepas *et al*. 2016).

### Gene expression quantification and differential expression analysis

Quantification of gene expression was performed using Kallisto v0.46.2 (Bray *et al*. 2016). The Kallisto index was built using all BRAKER2 predicted transcripts with the default k-mer size of 31. Transcripts per million (TPM) gene abundance values from Kallisto were scaled using the average transcript length, averaged over samples and to library size, using the lengthScaledTPM option in tximport v1.18.0 (Soneson *et al*. 2016). Quality control of the data was performed via principal component analysis, which showed clustering of all biological replicates and strong separation of crawling larva and juvenile slug sample types. Veliger and egg sample types also showed distinct clusters, but with slight overlap of one egg replicate close to 1.5 standard deviations from the veliger cluster in PC1 and PC2 space (Figure S7).

A differential expression analysis was performed to compare gene expression in the juvenile slug samples (post-feeding) to the three pre-feeding developmental stages (egg, veliger, and crawling larvae). Raw gene counts from Kallisto were passed to EdgeR v3.32.1 (Robinson *et al*. 2009), and only genes with an average TMP **>** 1 in at least one of the four developmental stages were retained. Raw gene counts were normalized using the TMM (trimmed mean of M values) method (Robinson and Oshlack 2010). Exact tests were conducted using a trended dispersion value and a double tail reject region. The false discovery rate (FDR) was calculated using the Benjamini-Hochberg (BH) procedure (Benjamini and Hochberg 1995).

### Identification and analysis of orthologous gene families

Homology between the predicted proteomes of *E. crispata* and four other gastropods (Table S5) was determined using OrthoFinder v2.5.4 (Emms and Kelly 2019) with sequence similarity searches performed by DIAMOND (Buchfink *et al*. 2015). When multiple protein isoforms were present, the longest protein sequence per gene was selected for the OrthoFinder analysis. Functional annotations were performed on the predicted proteome of *E. crispata* using the eggNOG-mapper v2 online service using default settings (Cantalapiedra *et al*. 2021). Hypergeometric tests for enrichment of functional categories were performed in python using the SciPy library hypergeom (Virtanen *et al*. 2020), and p-values were adjusted for multiple comparisons using the StatsModels library multitest with the BH method (Benjamini and Hochberg 1995; Seabold and Perktold 2010). GO terms were collapsed to medium length lists of representative GO terms using the simRel somantic similarity score in Revigo (Supek *et al*. 2011).

### PKS phylogeny

*EcPKS1* (Torres *et al*. 2020) was queried using blastp against the same custom database used for the Alien Index analysis, and full length sequences corresponding to the top 2000 significant hits (evalue < 1e^-10^) were extracted. All PKS homologs were queried using hmmscan (Eddy 2009) against three ketoacyl-synthase domain profiles (PF00108, Beta-ketoacyl synthase, N-terminal domain; PF00108, Beta-ketoacyl synthase, C-terminal domain; and PF16197, Ketoacyl-synthetase C-terminal extension), which were all present in *EcPKS1.* For each homolog, the maximum sequence region that spanned all three domains was extracted and retained for phylogenetic analysis; domain regions < 350 or > 650 amino acids were excluded. To reduce redundancy in the final phylogeny, database sequences were collapsed with CD-HIT v4.8.1 using a sequence identity threshold of 0.8 (Fu *et al*. 2012); sequences from kleptoplastic slugs were not collapsed. The final sequence set was aligned with MAFFT v7.471 using the L-INS-i strategy (Katoh and Standley 2013). Phylogenies were inferred using maximum likelihood as implemented in IQ-TREE v1.6.12 (Nguyen *et al*. 2015) using an empirically determined substitution model and 1000 rapid bootstrap replications. The phylogeny was visualized using ETE v3 (Huerta-Cepas *et al*. 2016).

### Data availability

All supplemental material as well as genome assembly, predicted CDS and protein sequences, multiple sequence alignments, tree files, and other related data files are available through FigShare (https://doi.org/10.6084/m9.figshare.23635812.v1). Scripts are available through GitHub (https://github.com/WisecaverLab/elysia_crispata_ECLA1_genome). *E. crispata* 28S ribosomal RNA gene sequence has been deposited at the NCBI GenBank database (http://www.ncbi.nlm.nih.gov/genbank) under accession OR177832. Raw sequencing reads used for *de novo* whole-genome assembly have been deposited in the Sequence Read Archive database under BioProject PRJNA987316.

## RESULTS

### *E. crispata* genome assembly and annotation

Sequencing data for the *E. crispata* genome assembly consisted of 18.98 Gbp of ONT long reads (2,504,236 reads with an N50 of 8,728 bp) and 75.25 Gbp of paired-end Illumina short reads. The Illumina data had an estimated heterozygosity of 0.88% with a projected genome size of 711.11 Mbp (Figure S8). The final genome assembly consisted of 8,089 contigs spanning 786.3 Mbp, with an N50 of 458.73 kbp. To evaluate completeness and coverage, we aligned the ONT and Illumina gDNA reads to the *E. crispata* genome assembly. Coverage histograms showed a single dominant peak at 20x and 86x coverage for ONT and Illumina reads, respectively (Figure S8), indicating that the genome assembly was largely homozygous without a significant amount of redundant haplotigs. The proportion of the genome with ≥10x read support was high: 751.15 Mbp (96.29%) using ONT reads and 757.17 Mbp (96.29%) using Illumina reads.

Repetitive elements made up 29.85% of assembly bases (Table S6). Most repetitive elements were unclassified, which comprised 17.7% of the genome assembly. Simple repeats made up 7.19%. of the genome. Classified retroelements and DNA transposons comprised 2.85% and 1.47% of the genome, respectively. The repeat content in *E. crispata* is comparable to the repeat content observed in the other sequenced saccoglossan genomes (29-33% repetitive) (Cai *et al*. 2019; Maeda *et al*. 2021).

A total of 68,514 genes encoding 71,367 transcripts were predicted using a combination of *ab initio*, homology-based, and transcriptome-based prediction methods (Table 1). The average protein-coding gene was 5,779.57 bp long and contained 4.82 exons. Functional annotations were ascribed to 24.23%, 15.51%, and 15.01% of genes using the PFAM, KEGG, and Gene Ontology databases, respectively (Table 1). Within the *E. crispata* protein-coding gene set, 96.65% (922/954) of BUSCO conserved metazoan genes were identified as complete, and of those, 94.03% were present in single-copy and 5.97% were duplicated (Table S3). Furthermore, 99.22% (253/255) of BUSCO conserved eukaryota genes were identified as complete, of those 92.50% were present in single-copy and 7.51% were duplicated (Table S3). This level of BUSCO recovery is the greatest of any currently available genome assembly from kleptoplastic sacoglossans (Figure 2A; Table S7).

**Figure 2.**
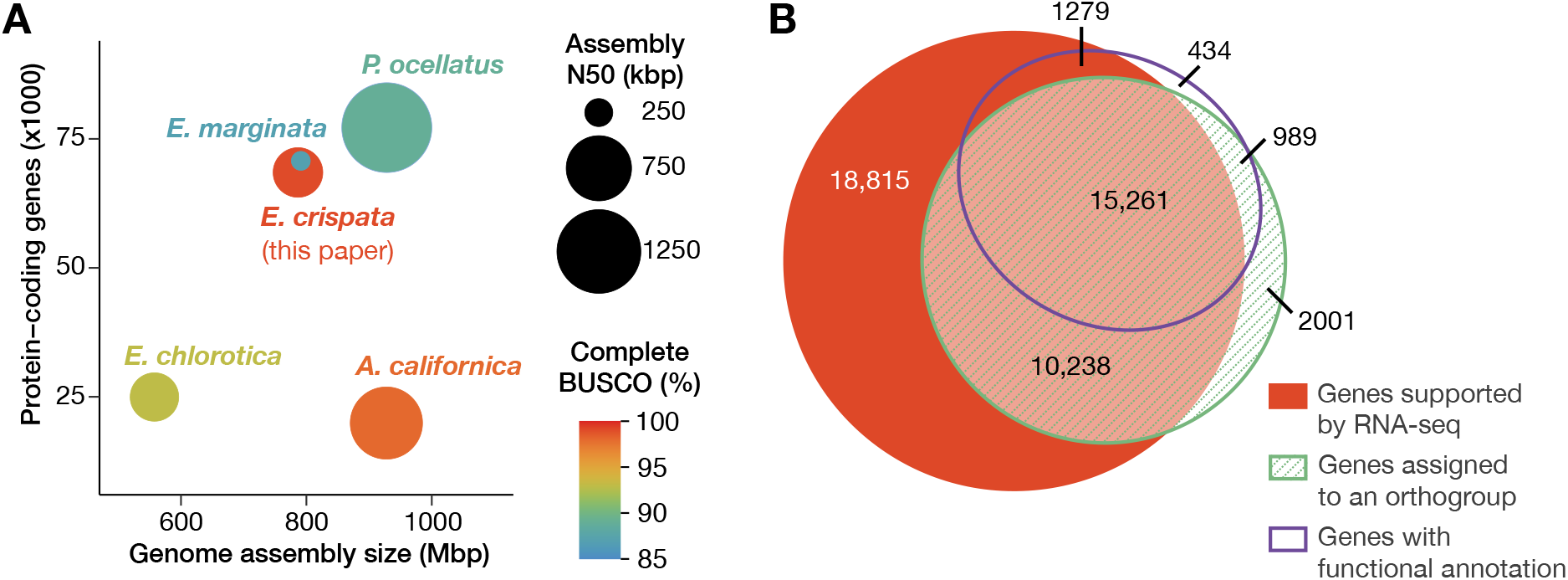
Genome assembly and annotation statistics. A) Comparison of assembly completeness and contiguity of four kleptoplastic sea slugs (*E. crisptata, E. chlorotica, P. ocellatus,* and *E. marginata*) and the sea hare *A. californica*. B) Area-proportional Venn diagram (Micallef and Rodgers 2014) depicting overlapping support for *E. crisptata* gene models. Support categories are as described in Table 1. Genes with two or more sources of support were assigned to the high-confidence gene set.

**Table 1.** Summary statistics of *E. crispata* gene models.

| Gene model statistics | Total gene set | High-confidence gene set |
| --- | --- | --- |
| No. protein-coding genes | 68,514 | 27,767 |
| No. transcripts | 71,367 | 30,043 |
| Mean gene length | 5,779.57 bp | 10,610.16 bp |
| Median gene length | 2,483 bp | 6,891 bp |
| Mean no. exons per transcript | 4.82 | 8.11 |
| Median no. exons per transcript | 3 | 5 |
| Mean exon length | 207.29 bp | 201.20 bp |
| Median exon length | 132 bp | 128 bp |
| No. genes supported by RNA-Seq<sup>1</sup> | 45,593 (66.55%) | 26,778 (96.44%) |
| No. genes with functional annotation<sup>2</sup> | 17,960 (26.21%) | 17,529 (63.13%) |
| No. genes assigned to an orthogroup<sup>3</sup> | 28,489 (41.58%) | 26,488 (95.39%) |
| No. genes with PFAM annotation | 16,601 (24.23%) | 16,220 (58.41%) |
| No. genes with GO annotation | 10,287 (15.01%) | 10,191 (36.70%) |
| No. genes with KEGG annotation | 10,624 (15.51%) | 10,498 (37.81%) |

<sup>1</sup>Average length-scaled TPM $\geq 1$ in at least one developmental stage: egg, veliger, larval slug, juvenile slug.
<sup>2</sup>Assigned one or more annotation via EggNOG-mapper.
<sup>3</sup>Excluding orthogroups unique to *E. crispata*.

The *E. crispata* genome assembly is similar to *E. marginata* in genome assembly size and number of protein-coding genes (Figure 2A). Gene count is noticeably elevated in *E. crispata, E. marginata,* and *P. ocellatus* (n > 68,000) compared to a fourth kleptoplastic Sacoglossa, *E. chlorotica* (n = 24,980), as well as the non-kleptoplastic sea hare, *Aplysia californica* (n = 19,945). We suspected the difference in gene count was due to an abundance of *de novo* predicted gene calls in the former species set. To address this, we determined the number of *E. crispata* predicted protein-coding genes with transcriptome- or homology-based support. Most of the *de novo* predicted genes in *E. crispata* (66.55%) showed evidence of expression, having an average length-scaled TPM > 1 in at least one of the four developmental stages surveyed (Figure 2B, Table 1). Fewer genes had homology-based support; 41.58% were assigned to gene families containing one or more species in addition to *E. crispata,* and 26.21% of genes were assigned one or more functional annotations via EggNOG-mapper (Figure 2B, Table 1). We created a high-confidence gene set of 27,767 genes, which we defined as those genes with two or more sources of support (*i.e.*, RNA-seq, orthology, and EggNOG-mapper). Median gene length and exons per transcript were increased in the high-confidence gene set compared to the total set (Table 1). Both gene sets can be downloaded from FigShare (see Data Availability). When accessing the data, we encourage users to select the gene set most appropriate for their biological question.

### Assessment of HGT

We calculated the Alien Index for all *E. crispata* predicted proteins to identify possible contamination and cases of HGT. The Alien Index screen flagged 194 contigs as bacterial contamination, which were subsequently excluded from the final assembly (see Materials and Methods). Only seven proteins were flagged as candidate HGTs with top hits to diverse organisms in Apicomplexa, Ochrophyta, Proteobacteria, and Riboviria (Table S8). None of the candidate HGTs had predicted functions associated with photosynthesis or plastids (Table S9). We used a custom phylogenetic pipeline to manually evaluate all candidate HGTs and found that all but one lacked phylogenetic support for HGT. The only HGT with phylogenetic support was Ecla3748g428790 (Figure 3), a single-exon gene with homology to a fragment of RNA-dependent RNA polymerase (RdRp) from negative-sense RNA viruses of invertebrates. The Ecla3748g428790 transcript had low but measurable expression support in our RNA-seq dataset with length-scaled TPM > 1 in three of four developmental stages surveyed (Table S9). RdRp genes appear to be frequently transferred into the nuclear genomes of their eukaryotic hosts (Dolja and Koonin 2018), supporting the inference that this sequence was horizontally acquired from a virus in *E. crispata*.

**Figure 3.**
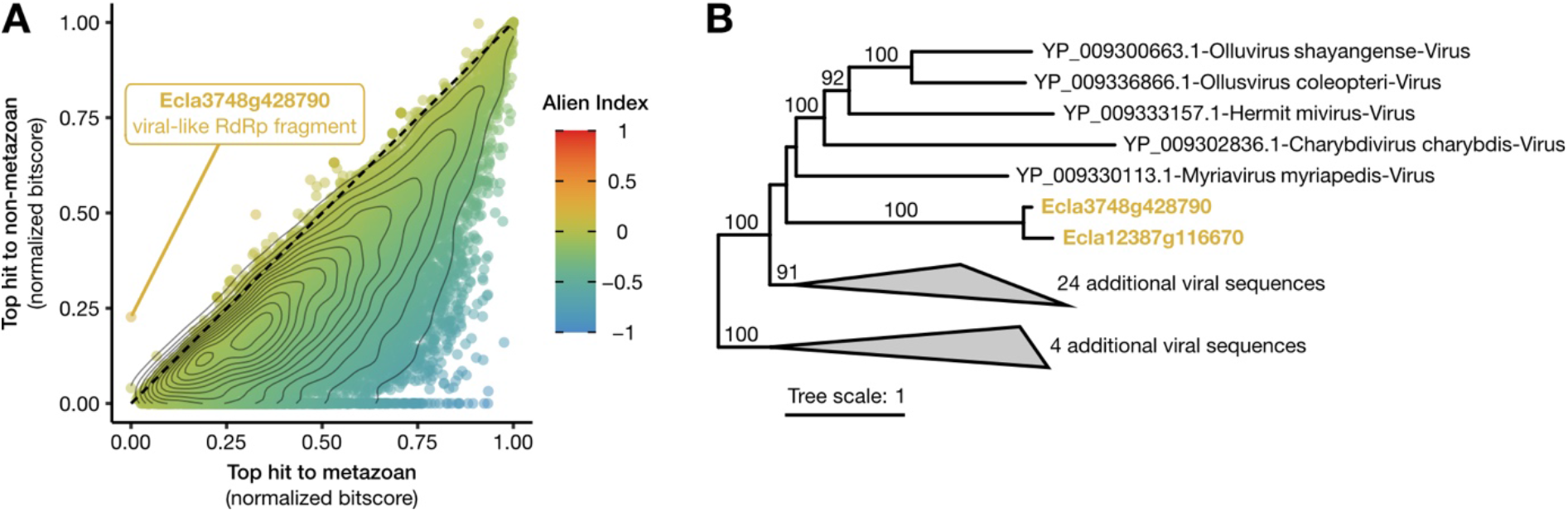
Detection of HGT in *E. crispata.* A) Alien Index analysis. Normalized bitscores for the top hit to the metazoan lineage (x-axis) are compared to the strength of the top hit outside of metazoans (y-axis). Genes above the dashed line have an Alien Index > 0 and are candidates for HGT. Contour lines indicate a 2D kernel density estimation of the distribution of Alien Index scores. Highlighted HGT candidate gene, Ecla3748g428790, has top hit to RNA-dependent RNA polymerase (RdRp) from Hubei myriapoda virus. No other HGT candidates were supported following phylogenetic analysis. B) RdRp phylogeny showing phylogenetic support for Ecla3748g428790 being horizontally acquired from viruses. Numbers along branches indicate ultrafast bootstrap support values (≥ 95) for descendant nodes.

### Phylogenetic placement

Slugs were identified as the clarki morphotype of *E. crispata* based on macroscopic morphological features, including a near-uniform green coloration with small round white spots, a transparent and tapered foot, and the presence of a gap in the parapodia in the mid-line near the head (Figure 1A,B,G) (Pierce *et al*. 2006; Krug *et al*. 2016). To confirm the placement of our assembled genome in the Sacoglossa species tree, we constructed concatenated maximum-likelihood phylogeny consisting of two nuclear loci coding for the 28S large ribosomal subunit and histone protein H3. Both 28S and H3 have served as DNA barcode genes in a previous phylogenetic analysis of the lineage (Händeler *et al*. 2009; Christa *et al*. 2014a, 2015; Krug *et al*. 2016; Martín-Hervás *et al*. 2021). The 28S sequence from the *E. crispata* genome was 99.58% identical (1406/1412 bp) to 28S amplified from another *E. crispata* collected from the Florida Keys (Genbank accn: KM230502, Krug *et al*. 2016), and the H3 genome sequence was 100% identical (327/327 bp) H3 amplified from the same sample (Genbank accn: KM040828, Krug *et al*. 2016), supporting its identification as *E. crispata* based on morphology. Genome assembly sequences grouped with *E. crispata* and sister species *E. ellenae* (Ortea *et al*. 2013; Krug *et al*. 2016) in both the concatenated and single-gene phylogenies with strong support (ultrafast bootstrap ≥ 98; Figure 4; Figure S6). Other *Elysia* species that grouped closely to *E. crispata* included *E. canguzua, E. chlorotica, E. diomedea, E. evelinae, E. serca,* and *E. viridis* (Figure 4), consistent with these species being members of a West Atlantic subclade complex (Krug *et al*. 2016).

**Figure 4.**
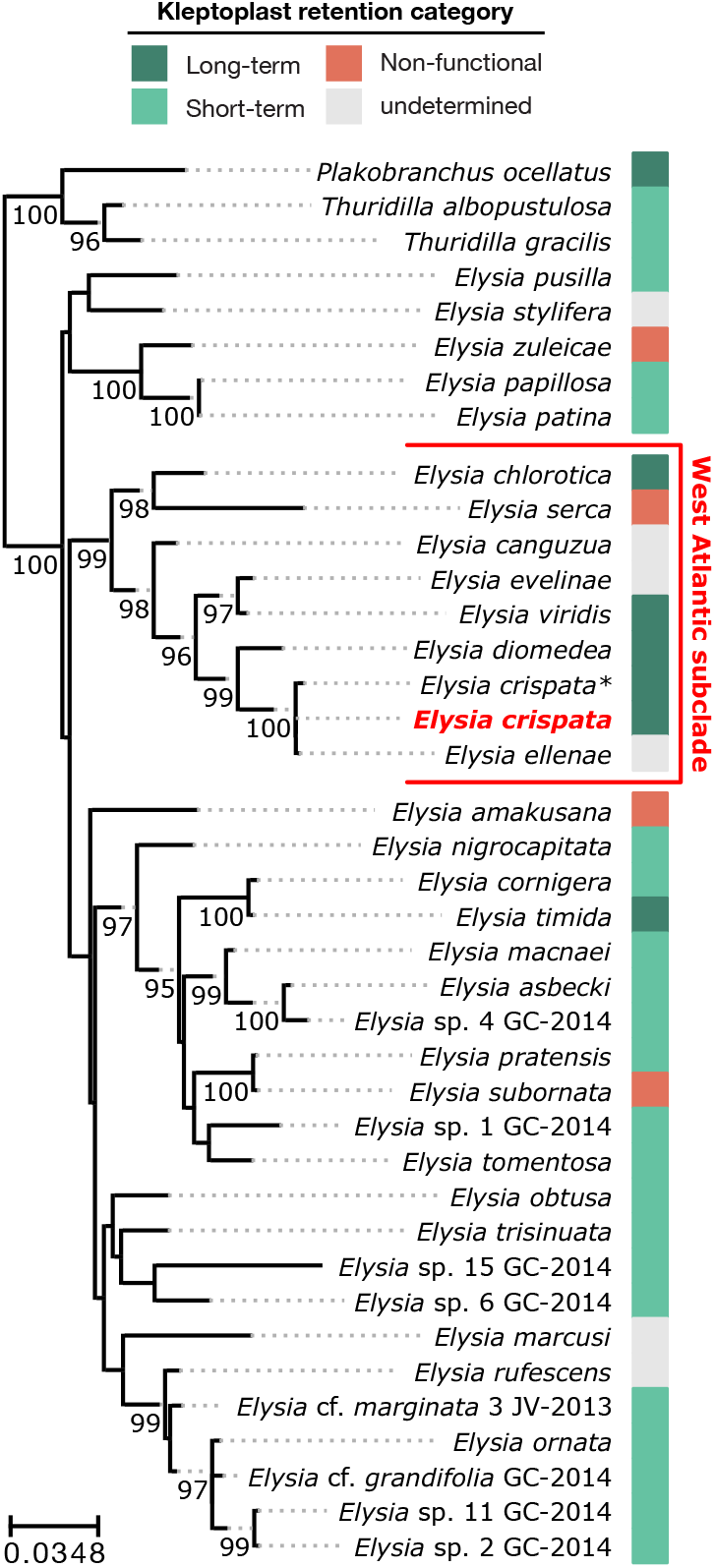
Phylogenetic analysis of sacoglossan sea slugs. Maximum likelihood analysis of concatenated 28S and H3 loci. *E. crispata* genome sequence is in red. Asterisk (*) indicates additional *E. crispata* sequence from Krug *et al*. (2016). Tree was rooted on the *Thuridilla/Plakobranchus* clade. Numbers along select branches indicate IQ-TREE ultrafast bootstrap support values (≥ 95) for the descendant nodes. The color bar indicates the kleptoplast retention state for each species: long-term retention (>20 days); short-term retention (>1 day and <20 days); see Table S12 for references. See Figure S6 for single-gene phylogenies of 28S and H3.

### Gene expression across developmental life stages

We tracked *E. crispata* gene expression across four developmental life stages. Three life stages (egg, veliger, and crawling larva) were grown in the absence of algae and contained no kleptoplasts. The fourth life stage, juvenile slug, was allowed to feed on macroalgae and acquired green pigmentation indicative of active kleptoplast sequestration (Figure 1G). Most predicted genes were not expressed under any of the four life stages (n = 22,925; average length-scaled TPM < 1); most genes in this category were not assigned to the high-confidence set (Figure 5A). In contrast, gene models that were expressed across all four developmental stages (n = 19,772; average length-scaled TPM > 1) were mostly comprised of high-confidence genes (Figure 5A). A total of 5,863 gene models were uniquely expressed in the kleptoplast-containing juvenile slugs with no measurable expression in the unfed life stages; of those, 2,044 gene models (34.9%) were considered high-confidence (Figure 5A).

**Figure 5.**
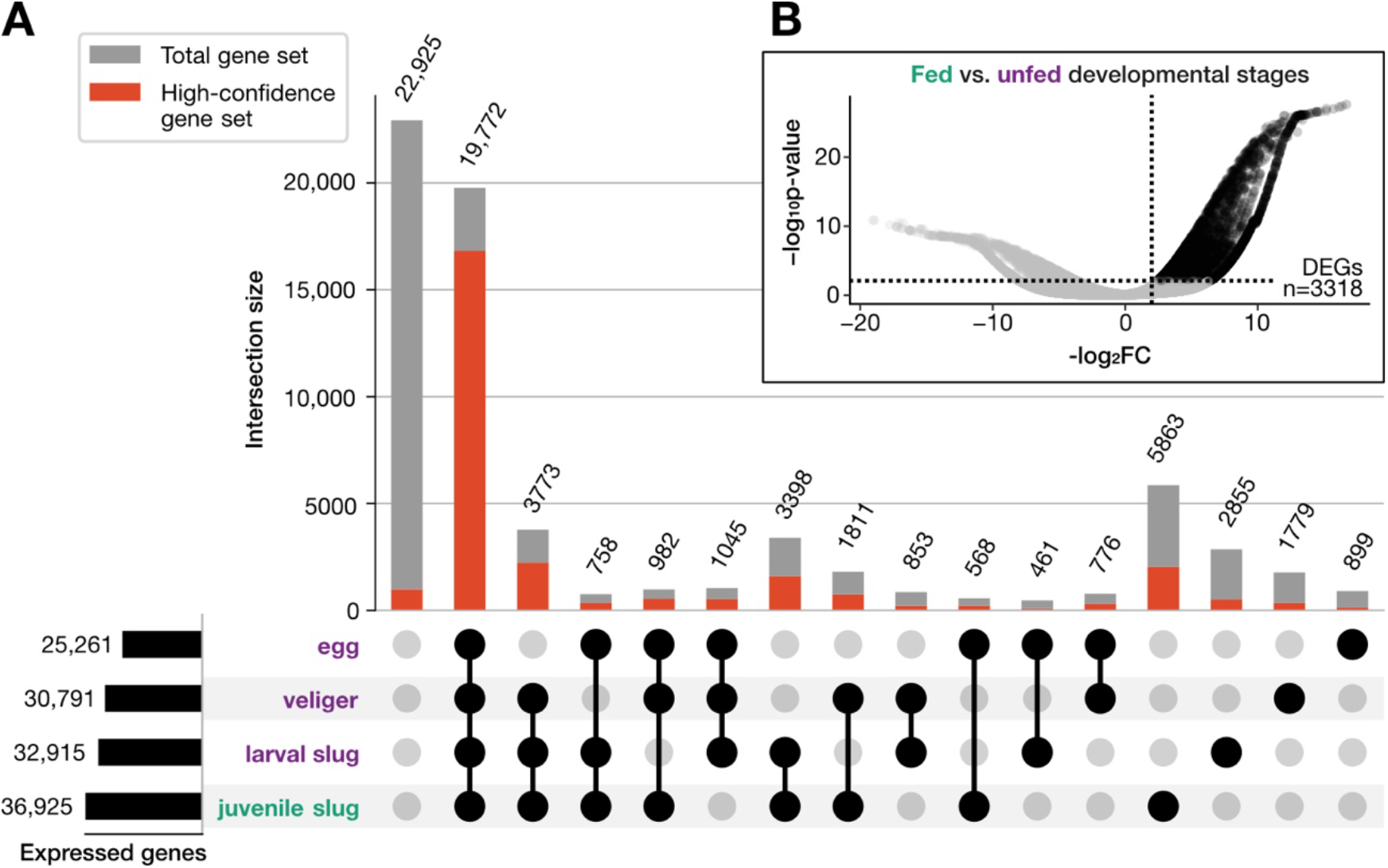
Gene expression in different slug life stages. A) Upset plot displaying intersection of gene expression across three unfed life stages (purple: eggs, veligers, and crawling larvae) and in algae-fed juvenile slugs (green). Filled circles and connecting lines indicate expression support under those conditions (average length-scaled TPM > 1). Numbers above grey bars indicate the total number of genes present in each intersection. Red bars indicate the proportion of each intersection comprised of high-confidence genes. B) Distribution of *E. crispata* gene expression comparing algae-fed juvenile slugs to unfed developmental stages. Dashed lines indicate significance thresholds for differential expression (-log_2_FC > 2; BH adjusted p-value < 0.01). Number indicates count of differentially expressed genes (DEGs) with increased abundance in fed slugs.

RNA-seq analysis identified 3318 differentially expressed genes (DEGs) that were significantly more abundant in kleptoplast-containing juvenile slugs compared to the unfed developmental life stages (-log_2_FC > 2; BH adjusted p-value < 0.01; Figure 5B). A test for significant enrichment of Gene Ontology (GO) functional categories in DEGs identified 561 enriched GO terms (hypergeometric test, BH-adjusted p-value < 0.05; Table S10). Among the most abundant were GO terms such as GO: 0051239, regulation of multicellular organismal process; GO:0032879, regulation of localization; GO:0009605, response to external stimulus; and GO:0002376, immune system process (Table S10). We hypothesize that DEGs in this analysis include candidate genes involved in long-term kleptoplast retention.

### Candidates for kleptoplasty-associated genes

To distinguish conserved gastropod developmental genes from genes potentially involved in kleptoplasty, we performed an OrthoFinder analysis (Emms and Kelly 2015) using the predicted proteome of *E. crispata* and four additional gastropod genomes (Figure 6). Included in our analysis were genomes from two sacoglossan slugs capable of long-term kleptoplast retention, *E. chlorotica* (Cai *et al*. 2019) and *P. ocellatus* (Maeda *et al*. 2021). A third Sacoglossa was capable of short-term kleptoplast retention, *E. marginata* (Maeda *et al*. 2021). Lastly, one non-kleptoplastic gastropod relative (the sea hare, *A. californica*) was also included (see Table S6 for genome accessions). The OrthoFinder analysis identified 119,336 orthogroups (predicted gene families), of which 21,821 were present in two or more species in the analysis. Excluding singletons, most orthogroups were present in all five gastropod species (n = 11,249; Figure 6A). Within these conserved orthogroups, 1,054 *E. crispata* homologs had significantly higher expression in juvenile slugs compared to the earlier developmental stages (Figure 6B). Functional enrichment of GO terms indicated that many of these genes in *E. crispata* have putative functions in the regulation of multicellular organismal processes (GO:0051239), developmental processes (GO:0032502, GO:0050793), and anatomical structure morphogenesis (GO:0009653) (Figure 6C; Table S10).

**Figure 6.**
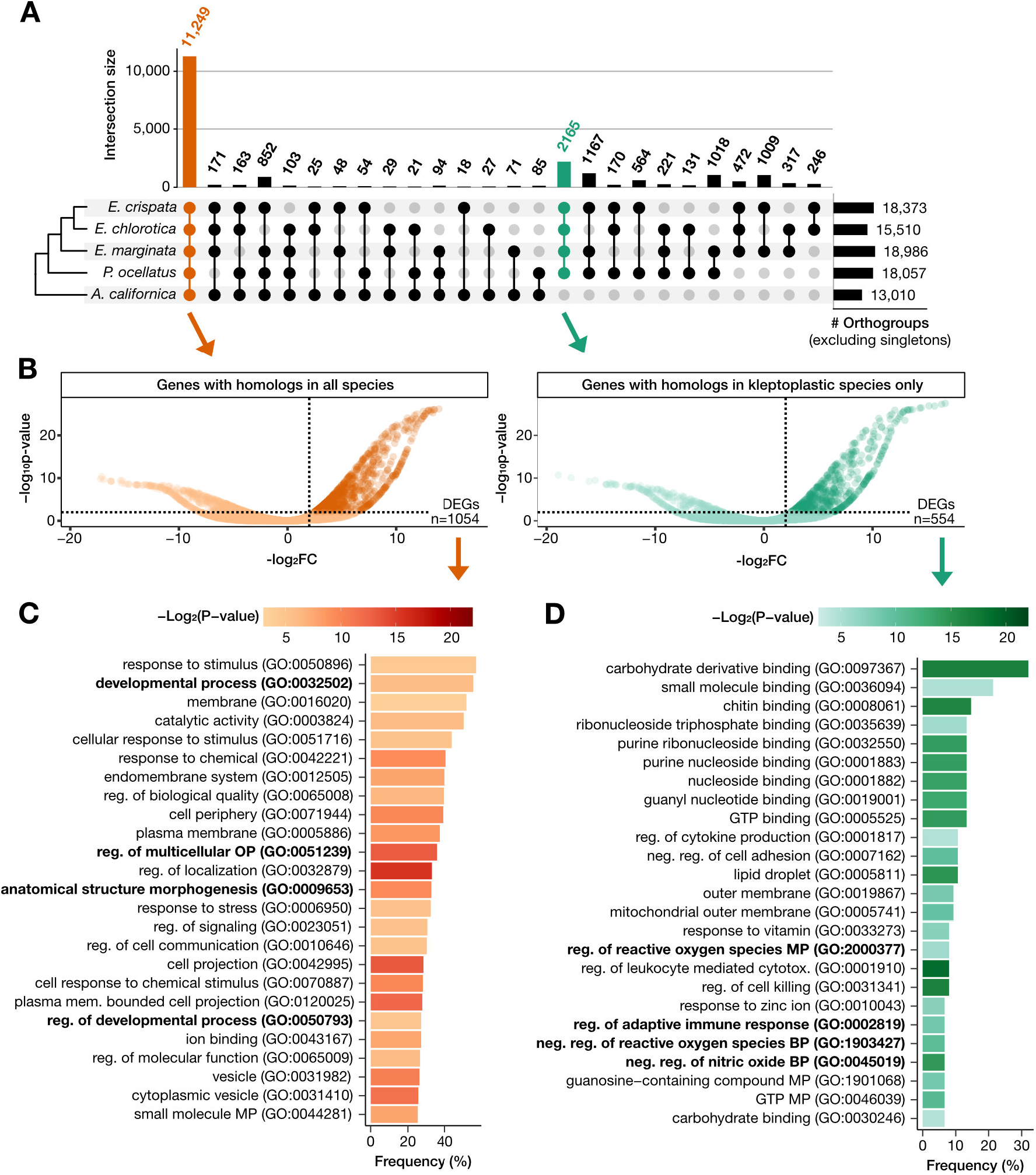
Gene family analysis of differentially expressed genes. A) Upset plot depicting intersection of orthogroups among four kleptoplastic sea slugs and the sea hare *A. californica*. Orange intersection indicates orthogroups present in all species; green intersection indicates orthogroups present in all kleptoplastic species and absent in the outgroup. B) For the orange and green intersections, differentially expressed genes (DEGs) with increased abundance in fed slugs compared to unfed life stages (see Figure 5B). Tests for enrichment of functional categories were performed on these two DEG categories: C) top 25 most abundant GO categories significantly enriched in DEGs with homologs in all other species; D) top 25 most abundant GO categories significantly enriched in DEGs with homologs in all other kleptoplastic species and absent in the outgroup (see Table S10). GO categories highlighted in the text are bolded.

A total of 2,165 orthogroups were present in all kleptoplastic Sacoglossa and absent in the non-kleptoplastic sea hare (Figure 6A). Within these sacoglossan-specific orthogroups, 554 *E. crispata* homologs had significantly higher expression in juvenile slugs compared to the earlier developmental stages (Figure 6B). These genes showed significant enrichment of GO categories involved in the regulation of reactive oxygen species (ROS) (GO:2000377, GO:1903427), regulation of nitric oxide (GO:0045019), and adaptive immune response (GO:0002819) (Figure 6D; Table S10).

Previous genomic and transcriptomic analyses of *P. ocellatus* identified three types of genes that were highly duplicated in the *P. ocellatus* genome and overexpressed in kleptoplast-containing tissues, suggesting they may be involved in kleptoplast retention (Maeda *et al*. 2021). We searched the *E. crispata* genome for homologs belonging to these three gene families: apolipoprotein D-like (KEGG: K03098), cathepsin D-like (KEGG: K01379), and lectin-like genes (KEGG: ko04091). The *E. crispata* genome contained eleven apolipoprotein D-like genes and six cathepsin D-like genes (Table S9), but neither gene family was enriched in genes with higher expression in kleptoplast-containing slugs compared to earlier developmental stages. In contrast, the *E. crispata* genome contained 307 lectin-like genes, 40.4% of which had higher expression in kleptoplast-containing slugs (Table S9). Moreover, lectins was the only KEGG functional category significantly enriched in the sacoglossan-specific gene set (hypergeometric test, BH-adjusted pvalue = 1.3e^-6^; Table S11).

A polyketide synthase in *E. chlorotica* (EcPKS1) was recently shown in vitro to synthesize the precursor for 7- and 8-propionate pyrones (Torres *et al*. 2020), which are proposed to function as UV- and oxidation-blocking compounds in slug cell membranes (Powell *et al*. 2018). *EcPKS1* and other type 1 iterative polyketide synthase genes in kleptoplastic slugs appear to have evolved from fatty acid synthase (FAS) genes present in all animals that are required for primary metabolism. We identified several PKS/FAS homologs in the *E. crispata* genome (Figure 7). One gene (Ecla2149g328170) grouped with FAS genes from *E. chlorotica* and other gastropods. Four genes in *E. crispata* were members of an expanded clade of sacoglossan-specific PKS-like genes (Figure 7B). Ecla12132g95200 (EclaPKS1) grouped with EcPKS1 in a subclade specific to species capable of long-term kleptoplast retention. Ecla7432g605620 (EclaPKS2) and Ecla7432g605630 (EclaPKS2-like) grouped with EcPKS2. In a third PKS clade, Ecla2359g352500 (EclaPKS3) grouped with an uncharacterized gene in *E. chlorotica* (Genbank accn: RUS77019), which we called EcPKS3 following the naming convention used previously by others (Torres *et al*. 2020). EclaPKS1 and EclaPKS3 were mostly highly expressed in kleptoplast-containing juvenile slugs (Figure 7C); however, neither were statistically significantly overexpressed in juvenile slugs compared to the unfed developmental stages (Table S9).

**Figure 7.**
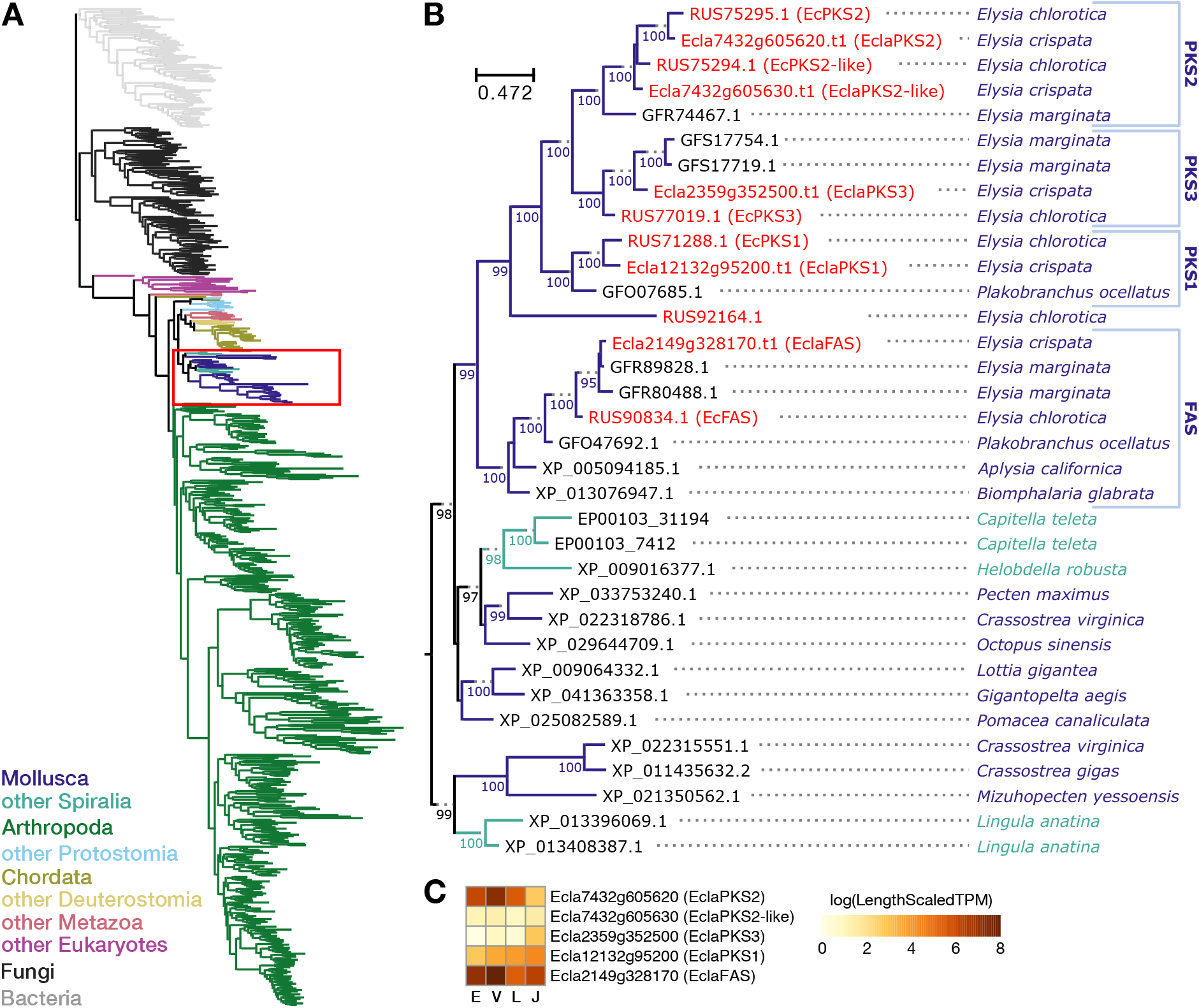
Phylogenetic analysis of FAS and PKS genes in kleptoplastic slugs. A) Maximum likelihood phylogeny of the ketosynthase domains with significant sequence similarity to the type 1 PKS sequence in *E. chlorotica* (EcPKS1). Tree is rooted on bacterial PKS sequences, and branches are color coded according to their taxonomic identity. Molluscan sequences are indicated by the red box. B) Detailed view of molluscan sequences. *E. crispata* and *E. chlorotica* sequences are in red. Numbers along branches indicate ultrafast bootstrap support values (≥ 95) for the descendant nodes. C) Average gene expression for *E. crispata* FAS/PKS homologs in four developmental stages (E, egg; V, veliger; L, larval slug; J, juvenile slug).

## DISCUSSION

The *E. crispata* genome assembly serves as a high-quality reference for future functional and evolutionary studies in kleptoplastic sea slugs. With calculated BUSCO scores of 93-99% (Table S7), the *E. crispata* gene space is the most complete of all sequenced sacoglossans (Figure 2) and contends with the most complete of all mollusk genomes (Caurcel *et al*. 2021). As in previous genome-level analyses of kleptoplastic sea slugs (Wägele *et al*. 2011; Bhattacharya *et al*. 2013; Maeda *et al*. 2021), we found no evidence of HGT from host algae. Nevertheless, our search strategy detected one case of probable HGT from viruses (Figure 3). Alternative mechanisms for kleptoplast maintenance must be considered and investigated to better understand how these sea slugs can retain functional kleptoplasts for so long in the absence of algal nuclear-encoded chloroplast genes.

The ability to retain long-term kleptoplasts has evolved multiple times independently in Saccoglossa, but the exact number and timing of transitions from short-term to long-term kleptoplasty remains unclear (Christa *et al*. 2015). *E. crispata* belongs to the *Elysia* west Atlantic subclade (Krug *et al*. 2016), a lineage that is unique in that it contains several species that retain long-term kleptoplasts including *E. crispata, E. chlorotica, E. diamedea,* and *E. viridis* (Figure 4; Table S12). The prevalence of long-term kleptoplasts in this subclade suggests that long-term retention may be an ancestral characteristic of this lineage. However, another species within the subclade, *E. serca,* does not retain functional kleptoplasts (Clark *et al*. 1990), and several members of this subclade (e.g., *E. canguzua, E. ellenae,* and *E. evelinae*) are yet to be studied for kleptoplast retention. Characterizing the retention time for these species and determining the timing of transition(s) to long-term kleptoplasty in the west Atlantic subclade would inform our understanding of how kleptoplasty evolved in sacoglossan slugs. A single ancestral origin for long-term retention in this subclade would suggest that one or more rare innovations facilitated this key transition. In contrast, if long-term retention arose repeatedly in the subclade (*i.e.,* parallel or convergent evolution), the transition to long-term retention may be the result of relatively simple genetic mechanisms. Addressing this question requires additional taxonomic sampling and phenotyping in this group.

Combined, comparative genomic and gene expression analyses identified 554 genes in *E. crispata* that were both unique to kleptoplast-containing slugs and more highly expressed in juvenile slugs compared to unfed early developmental stages. Genes within this intersection included lectin carbohydrate-binding proteins (Table S11) as well as those involved in regulation of ROS, production of nitric oxide, and adaptive immunity (Figure 6D). Similar gene families were identified in a genomic analysis of *P. ocellatus,* another long-term kleptoplastic Sacoglossa (Maeda *et al*. 2021), providing additional evidence that genes in these categories may play a role in prolonged kleptoplast retention in these slugs. Recently, Torres *et al*. (2020) identified three type 1 polyketide synthase genes in *E. chlorotica* that may play a role in protecting the kleptoplast from UV and oxidative damage. Our analysis of this gene family identified four PKS genes in *E. crispata* (Figure 7), including one belonging to a new PKS3 subclade, suggesting that polyketide-based products in kleptoplastic slugs may be more diverse that previously recognized. Future work is necessary to develop *E. crispata* into a genome-enabled model system for functional investigation of these genes and others to better understand the genetic and cellular mechanisms of kleptoplasty.

## ACKNOWLEDGEMENTS

We thank members of the Wisecaver and Widhalm labs for helpful discussions. We also thank Dr. Dave Sandstrom (University of Maryland) for guidance on sea slug rearing and culturing algae. This work was conducted in part using the resources of the Rosen Center for Advanced Computing at Purdue University. This work was supported by Purdue University through research seed grants provided by the Center for Plant Biology and the Agricultural Science and Extension for Economic Development to JRW and JHW. This work was also supported by the National Science Foundation under grant DEB-1831493 to JHW, and by Research Corporation for Science Advancement award number 26212, by Showalter Research Trust award 41000747, and by Gordon and Betty Moore Foundation award 9331 to JRW. This work was also supported by the USDA National Institute of Food and Agriculture Hatch Project numbers 177845 to JRW and 1016057 to JHW.

